# Somatomotor Beta Bursts Mediate the Negative Impact of PTSD Severity on Conflict Monitoring

**DOI:** 10.1101/2022.12.23.521828

**Authors:** Eric Rawls, Craig A. Marquardt, Scott R. Sponheim

## Abstract

Cognitive control deficits are associated with posttraumatic stress disorder (PTSD) and may explain how reminders of past traumatic events intrude upon daily experiences of people who have experienced trauma. Lateralized somatomotor beta-band desynchronization, an electrophysiological signature of controlled movement, indexes the downstream output of cognitive control processes. Recent evidence suggests that somatomotor beta activity does not manifest as rhythmic oscillations, but instead as discrete and stochastic burst-like events. Here, we quantified the rates of lateralized somatomotor beta bursts (beta burst rates; BBR) evoked during a flanker cognitive control paradigm among United States military veterans from Operations Iraqi and Enduring Freedom (OEF/OIF) who show varying degrees of PTSD. We found BBR reflected both response direction and conflict monitoring during processing of stimuli that evoked response conflict. Impaired behavioral performance and increased peri-response BBR were related to greater posttraumatic stress symptomatology (PTSS). Critically, increased BBR mediated the link between PTSS and decreased conflict monitoring accuracy. Results suggest that poor cognitive control in PTSS reflects a failure to adaptively disinhibit target motor representations, rather than a failure to inhibit distractor representations. Thus, BBR reveal limited representation of target stimuli as a primary contributor to impaired cognitive control in PTSD. Because BBR were robustly associated with behavioral performance and exhibited high statistical reliability the index may carry utility for appraising individual differences in cognitive control in other brain disorders.

## 1 Introduction

Posttraumatic stress disorder (PTSD) is a debilitating condition that will impact ∼8% of the US population within their lifetime (Kessler et al., 2005, 2012), and as many as 15% of US military veterans returning from deployments during Operations Enduring Freedom or Iraqi Freedom (Tanielian et al., 2008). PTSD is characterized by recurring intrusive thoughts of traumatic events, maladaptive avoidance, negative changes in affect or mood, and hyperarousal (American Psychiatric Association, 2013). Deficits in cognitive control, or the ability to control our actions in pursuit of long-term goals (Botvinick et al., 2001), are also apparent in PTSD. Cognitive control is particularly relevant to the ability to suppress intrusive thoughts (Bomyea & Lang, 2015, 2016; Brewin & Beaton, 2002), a key feature of PTSD.

Cognitive control can be assessed using laboratory tasks that simultaneously prime mutually incompatible responses, such as the Flanker (Eriksen & Eriksen, 1974) and Stroop (Stroop, 1935) paradigms. Such tasks engage conflict monitoring, a subdomain of cognitive control that involves detecting when incompatible response patterns are activated and arbitrating between them to execute the correct response (Botvinick et al., 2001). This monitoring function relies on interactions between anterior cingulate, prefrontal, and premotor/motor cortices (Botvinick, 2007; Botvinick et al., 2004; Gehring & Knight, 2000), a neural circuit that contributes to our ability to regulate emotional responses (Ochsner & Gross, 2005). In this framework, ACC is thought to assume the role of monitoring for response conflict, and DLPFC is thought to take the role of implementing control. Premotor/motor cortex is densely connected with the ACC and DLPFC (Botvinick et al., 2001), and receives these downstream control signals to dynamically adjust motor inhibition. In the present investigation we sought to better understand neural processes related to conflict monitoring that might account for impaired cognitive control in posttraumatic psychopathology.

Cognitive control processes in prefrontal brain regions are associated with theta-band (4-8 Hz) activity (Cavanagh et al., 2012; Cavanagh & Frank, 2014). However, motor signals representing the output of conflict monitoring systems, and corresponding to directed movement, are most prominent in the beta frequency band (15-30 Hz). Motor cortical activity is sensitive to competition among choice stimuli prior to choice commission (Cisek & Kalaska, 2005; Pape & Siegel, 2016; Pastor-Bernier & Cisek, 2011), and predicts eventual choice during perceptual decision-making (Donner et al., 2009).

Lateralized somatomotor beta power also reflects cognitive control influences over motor output. Fischer et al. (2018) demonstrated that motor beta reflects post-error regulation in the flanker task, and specifically appears to gate the rate at which environmental information informs responding. As such, motor cortical signaling furnishes a readout of the downstream influence of conflict monitoring, potentially providing a neural indicator of clinically significant impairments in cognitive control.

Investigations of activity from non-invasive electrophysiological sensors over the human motor cortex have shown that power decreases in a broad beta-frequency (15-30) Hz band during the several hundred milliseconds that surround a behavioral response. However, investigations of beta-frequency neural activity in unaveraged single-trial traces suggest that beta activity is not rhythmic, but instead is better characterized as a series of transient burst events, each lasting approximately 150 ms (Lundqvist et al., 2016; Sherman et al., 2016). In primary somatosensory cortex, differences in pre-stimulus beta power that predicted detection of tactile stimuli (Jones et al., 2010) were later shown to instead reflect a difference in the rate of averaged pre-stimulus beta burst events (Shin et al., 2017), and the rate of beta burst events measured from somatomotor EEG sensors predicted response commission and successful response inhibition more strongly than averaged beta power (Wessel, 2020).

In the present study we examined conflict monitoring and posttraumatic psychopathology in 130 US military veterans who were previously deployed to combat zones. We hypothesized that the rate of somatomotor beta burst events would reflect conflict monitoring by signaling response direction and stimulus congruency in a flanker paradigm. Specifically, since beta burst rate decreases are thought to reflect disinhibition of motor cortex during movement, we hypothesized that beta burst rates would decrease over motor cortex contralateral to the responding hand for both congruent and incongruent trials. Furthermore, we hypothesized that ipsilateral beta burst rates would be lower for incongruent trials (compared to congruent), reflecting disinhibition of ipsilateral motor cortex by distracting flanker stimuli. We then investigated whether beta burst rates during conflict monitoring might characterize PTSD-relevant impairments in cognitive control. Critically, increased rates of peri-response beta burst events were systematically related to increased severity of posttraumatic stress symptomatology, and statistically mediated the link between PTSD symptom severity and impaired conflict monitoring.

## 2 Results

### 2.1 Lateralized Response Conflict Paradigm Demonstrates Cognitive Control Deficits in PTSD

Participants showed the lateralized response conflict effects typical for behavior on the flanker task. The mean accuracy rate was 90.9±7.7% (congruent: 96.3±4.6%, incongruent: 85.7±10.7%). Mean response times (RTs) were 455±44 ms (congruent: 409±40 ms, incongruent: 501±48 ms). A linear mixed-effects model (LMM) on accuracy rates demonstrated the expected within-subject effect of stimulus congruency, Wald χ2(1) = 137.30, *p* < .001, but no effect of group (all *p* > .1). Results were equivalent whether or not we included individuals with subthreshold PTSD diagnoses in the PTSD diagnostic groups. However, a model that considered overall PTSD symptom severity instead of PTSD diagnosis showed a main effect of symptom severity, Wald χ2(1) = 5.05, *p* = .025 (*Figure 1B*). This effect was equivalent when we instead used a robust model to predict accuracy rates, *t*(127.99) = -2.35, *p* = .020. A model that instead considered individual symptom subscales from the CAPS (instead of overall symptom severity) revealed a main effect of Intrusion symptom severity on accuracy, Wald χ2(1) = 4.98, *p* = .026, indicating lower accuracy with increasing intrusion symptom severity (*Figure 1C*). This effect was equivalent when we instead used a robust model to predict accuracy rates, *t*(123.99) = -2.57, *p* = .011.

**Figure 1.**
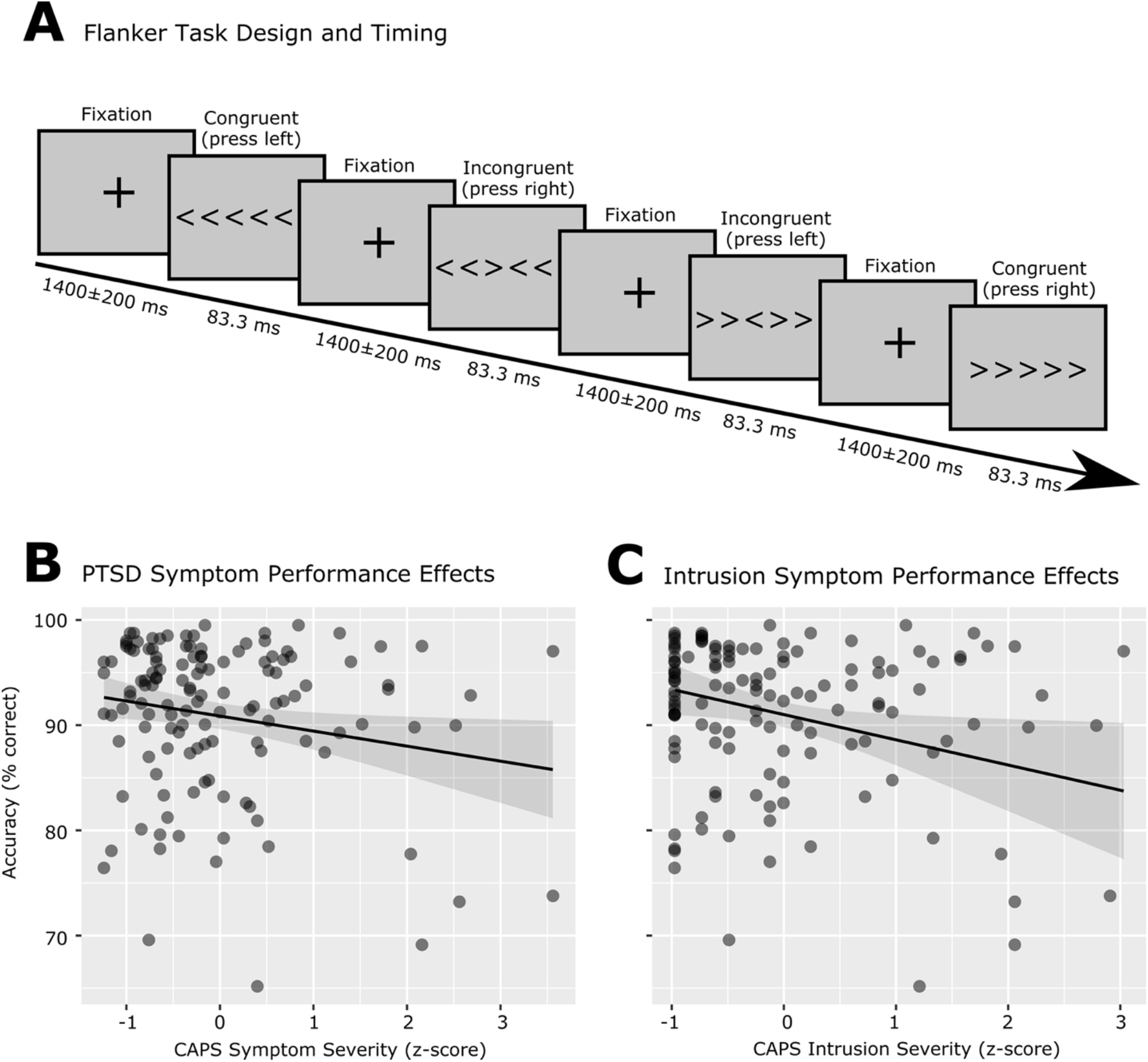
Lateralized Response Conflict Paradigm Reveals Lower Cognitive Control Related to Worse PTSD Symptomatology. A: Participants (n = 130) completed a flanker task while we recorded dense-array EEG. B: Worse task accuracy was predicted by increased overall PTSD (i.e., CAPS) symptom severity (p = .025). C: Worse task accuracy was predicted by increased intrusive reexperiencing of traumatic events (i.e., Intrusion) severity (p = .026).

This model also indicated a significant main effect of blast mTBI severity, Wald χ2(1) = 7.47, *p* = .006, indicating increasing accuracy with increasing blast mTBI severity. This effect was equivalent when we instead used a robust model to predict accuracy rates, *t*(123.99) = 3.09, *p* = .002. RTs demonstrated the expected within-subject effect of congruency, Wald χ2(1) = 939.59, *p* < .001, but no effects of diagnostic group or of symptom severity. As such, our analyses reveal response control deficits associated with overall PTSD symptom severity, but not a formal diagnosis of PTSD, as well as likely confounded response control benefits associated with increasing blast mTBI severity.

### 2.2 Somatomotor Beta Burst Rates Track Competing Motor Representations During Conflict

We examined the motor cortical outputs of conflict monitoring processes by detecting beta-frequency (15-29 Hz) bursts in single trials of cleaned EEG. Grand-average topographical plots of BBR are presented in *Figure 2A*. Single trials of beta-band activity at lateral somatomotor sensors clearly indicated the presence of burst-like activity that was brief (∼150 ms) and varying in time and frequency over trials (*Figure 2B*).

**Figure 2.**
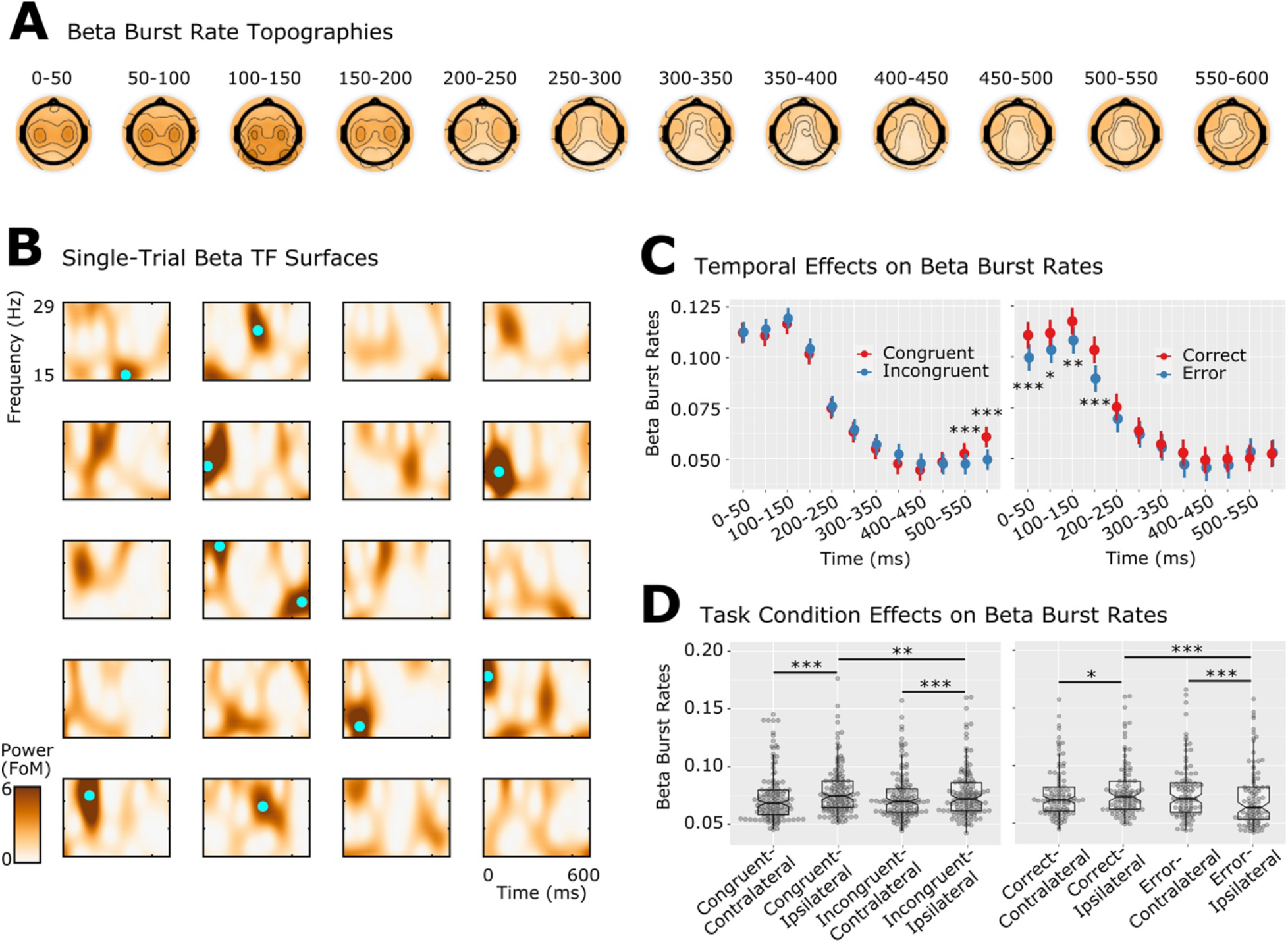
Somatomotor beta burst rates (BBR) track competing motor representations during conflict. A: Topographies of BBR, averaged within 50 ms windows (all conditions, 0 ms is stimulus onset). B: Example time-frequency surfaces of C3/C4 beta-band activity in single trials for randomly selected single subjects (one subject per row). Note that beta activity is brief and varying in time across trials (i.e., burst like). Detected bursts are superimposed on TF surfaces as cyan dots. Color scaling is in Factors of Median (FoM) as data are normalized according to frequency-specific medians. C: Left: Time Window X Congruency effects on averaged BBR. As described in Section 4.6 (Methods), the axis is interpreted as the average number of beta bursts in each time window per trial. BBR were increased for congruent compared to incongruent trials, but only for late time windows. Right: Time Window X Accuracy effects on averaged BBR. BBR were increased for correct compared to error trials, but only for early time windows. * p < .05, ** p < .01, *** p < .001. D: Left: Congruency X Laterality effects on averaged BBR. As described in Section 4.6 (Methods), the axis is interpreted as the average number of beta bursts in each time window per trial. BBR were reduced for contralateral compared to ipsilateral sensors indicative of less motor inhibition, for both congruent and incongruent stimuli. BBR were also reduced for incongruent compared to congruent trials, but only for ipsilateral sensors. Right: Accuracy X Laterality effects on averaged BBR. BBR were reduced for error trials compared to correct trials at sensors ipsilateral to the target, indicating increased motor engagement with the distractor stimuli during erroneous responses. Meanwhile, Laterality effects within correct/error trials both indicated that BBR were reduced contralateral to the response direction. * p < .05, ** p < .01, *** p < .001.

We first characterized somatomotor beta bursts in correct trials using a LMM containing within-subject factors of Congruency, Laterality, and Time Window. This analysis revealed main effects of Time Window, Wald χ2(11) = 9679.18, p < .001, and of Laterality, Wald χ2(1) = 54.76, *p* < .001, as well as interactions between Laterality and Congruency, Wald χ2(1) = 4.71, *p* = .030, and between Congruency and Time Window, Wald χ2(11) = 56.99, *p* < .001. The effect of Time Window indicated that beta burst rates decreased steadily from ∼150 ms post-stimulus up to the time of response (∼400-500 ms), followed by a significant post-response rebound. The interaction of Congruency and Time Window demonstrated that in late response windows, incongruent trials have lower BBR than congruent trials (time: 500-600 ms, |z| > 3.36, *p* < .001; *Figure 2C, left panel*). These results remained significant when *p*-values were corrected for n=12 multiple comparisons via the FDR (Benjamini & Hochberg, 1995) method. This likely represents slower motor processing induced by distractor stimuli, which activate conflicting motor representations and slow correct response activation. Finally, the interaction of Congruency and Laterality revealed that the effect of Laterality was significant in both congruent and incongruent trials, with lower BBR for contralateral, compared to ipsilateral, sensors (|z| > 3.69, *p* < .001; *Figure 2D, left panel*). The effect of Congruency was significant over ipsilateral sensors (z = 2.64, *p* = .008), with higher burst rates for congruent trials compared to incongruent trials, but there was no influence of Congruency over contralateral sensors (z = -0.41, *p* > .6; *Figure 2D, left panel*). Thus, we interpret this effect as most likely reflecting disinhibition of competing motor responses by distracting flanker stimuli.

Next, we characterized somatomotor beta bursts in incongruent trials using within-subject factors of Outcome, Laterality, and Time Window (*Figure 2C, right panel*). This analysis revealed main effects of Outcome, Wald χ2(1) = 29.47, *p* < .001 and of Time Window, Wald χ2(11) = 2863.36, *p* < .001, as well as interactions between Outcome and Laterality, Wald χ2(1) = 20.23, *p* < .001, and between Outcome and Time Window, Wald χ2(11) = 26.57, *p* = .005. Interpretation of the interaction of Outcome and Time Window demonstrated that BBR were significantly higher for correct trials (compared to error) during pre-response periods consisting of 0-200 ms (|*z*| > 2.54, *p* < .01; *Figure 2C, right panel*). These results remained significant when *p*-values were corrected for n=12 multiple comparisons via the FDR (Benjamini & Hochberg, 1995) method. Interpretation of the interaction of Outcome and Laterality demonstrated that correct trials had significantly higher BBR than error trials at sensors ipsilateral to the flanker target stimulus (representing processing of distractors), *z* = 7.02, *p* < .001, but not at sensors contralateral to the flanker target stimulus (representing processing of targets), *z* = 0.65, *p* > .5 (*Figure 2D, right panel*). Thus, incongruent trial errors are primarily associated with decreased inhibition at ipsilateral somatomotor sites, representing increased engagement of distractor motor responses.

### 2.3 Somatomotor Beta Burst Rates are Reliable and are Associated with Individual Differences in Response Control

We found that correct-trial BBRs had good split-half reliability (r > .7) during early time periods, and reliability increased to become very good peri-response (r > .8; *Figure 3A*). Error-trial BBRs had notably lower and more variable split-half reliability estimates, but still had adequate reliability peri-response (r > .65; *Figure 3A*). As such, we consider the psychometric properties of BBRs to be well-suited for individual differences analyses. We further observed that individual differences in accuracy rates negatively correlated with correct-trial BBR in time windows corresponding to 250-550 ms for congruent trials, and in time windows corresponding to 300-600 ms post-stimulus for incongruent trials (*Figure 3B*). These correlation coefficients were similar across ipsilateral and contralateral sensors. This is interesting, as one might expect the directions of these correlations to be in opposite directions for contralateral and ipsilateral sensors (at least for incongruent trials). We suggest this might be due to the high correlation (r > .9) between contralateral and ipsilateral beta burst rates, such that subjects with high ipsilateral BBR almost always had high contralateral BBR. Meanwhile, we observed that individual differences in response times positively correlated with BBRs in the pre-response period, with minor differences between conditions (*Figure 3C*). Error-trial BBR did not systematically relate to individual differences in behavior. As such, these results replicate and extend those of (Wessel, 2020), who demonstrated positive relationships between BBR and response times in a stop-signal task.

**Figure 3.**
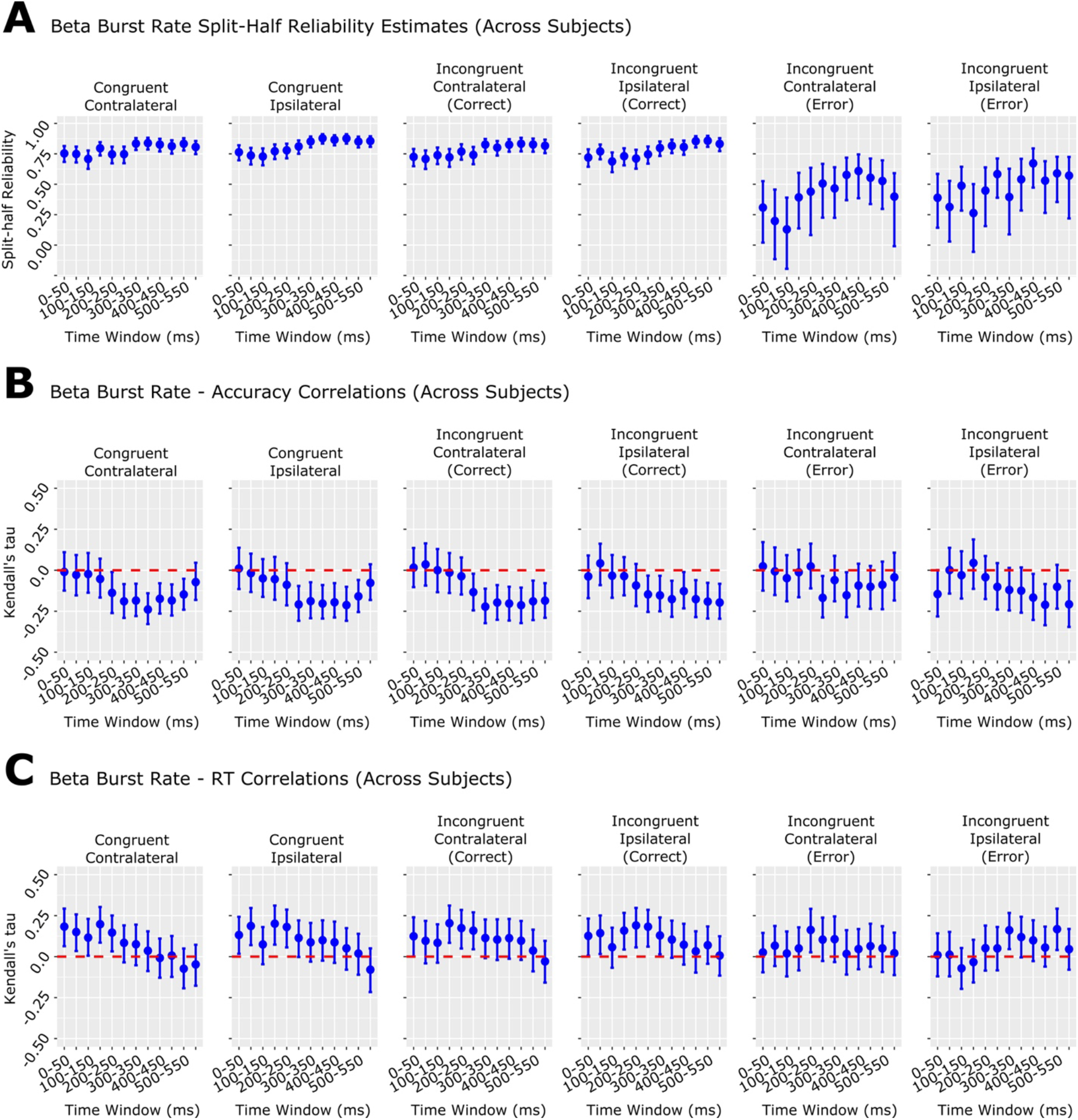
Somatomotor beta burst rates are reliable and are associated with individual differences in conflict monitoring. A: Across-subject beta burst rate (BBR) split-half trial-averaged reliabilities (Pearson correlations, Spearman-Brown corrected; bars = 95% CI). Split-half reliabilities were estimated by randomly generating two split-half averages per subject (n = 130 averages per vector; 5000 repetitions) and correlating these estimates across subjects. B: Across-subject trial-averaged BBR correlations with flanker task accuracy (ACC; Kendall’s tau; bars = bootstrap 95% CI). C: Across-subject trial-averaged BBR correlations with flanker task reaction times (RT; Kendall’s tau; bars = bootstrap 95% CI).

### 2.4 Correct-Trial Somatomotor Beta Burst Rates Are Related to Overall PTSD Symptom Severity

Thus far we have established that BBR track flanker task experimental manipulations including lateralized response direction and stimulus congruency. We next asked whether BBR might provide a neurobiological marker of cognitive control dysfunction after military deployment for individuals with PTSD symptomatology. The addition of diagnostic groups (controls/PTSD/mTBI/PTSD+mTBI) to our previous analysis of correct-trial BBR demonstrated an interaction of diagnostic group by time window, Wald χ2(33) = 92.42, *p* < .001, but post-hoc interpretation of this effect found no significant group differences in BBR in any time window (all *p* > .2). Results of this group-based analysis were equivalent when subjects with subthreshold PTSD were included in the PTSD groups.

A model that included overall PTSD symptom severity (as assessed via the CAPS) rather than a formal diagnosis of PTSD showed an interaction between PTSD symptom severity and time window, Wald χ2(11) = 51.67, *p* < .001. Post-hoc examination of the linear trend of CAPS within each time bin indicated that PTSD symptom severity predicted increased BBR in time windows corresponding to 250-300 ms (z = 2.07, *p* = .039), 300-350 ms (z = 2.31, *p* = .021), 350-400 ms (z = 2.35, *p* = .019), 450-500 ms (z = 2.35, *p* = .019), and 500-550 ms (z = 1.98, *p* = .047; *Figure 4A)*. These results fell below significance (all *p* > .08) when *p*-values were corrected for n=12 multiple comparisons via the FDR (Benjamini & Hochberg, 1995) method.

**Figure 4.**
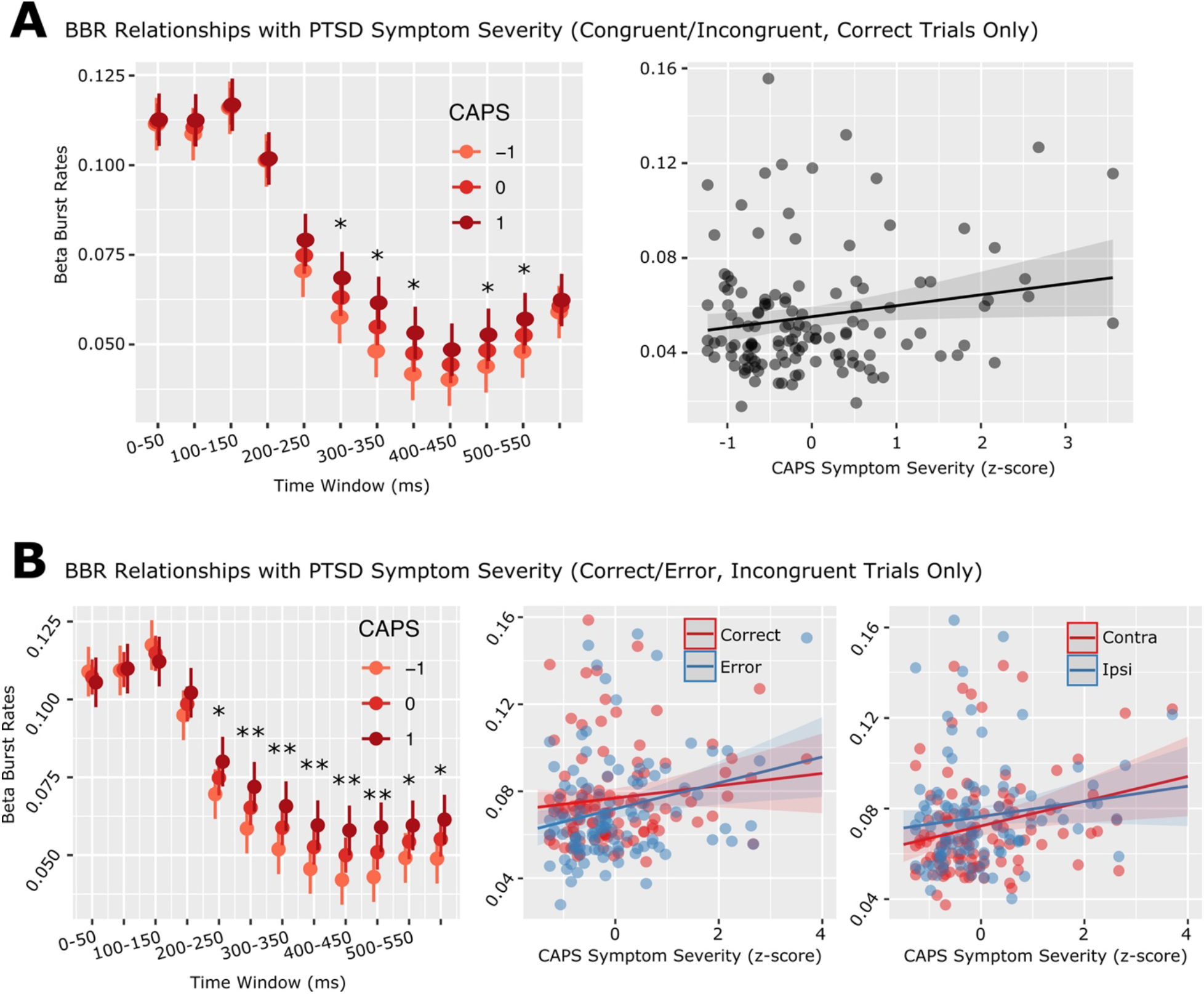
Somatomotor Beta Burst Rates Are Systematically Related to Overall PTSD Symptom Severity. A: Overall PTSD symptom severity (CAPS; shown at z-scores of -1, 0, and 1) effects on correct-trial BBR by time window (left panel; * uncorrected p < .05) and averaged over 250 to 550 ms time windows (right panel). Note that these effects fell below significance when results were corrected for n=12 comparisons (p > .08). B: Overall PTSD symptom severity (CAPS; shown at z-scores of -1, 0, and 1) on incongruent-trial BBR. Left panel: results by time window (* uncorrected p < .05, ** uncorrected p < .01). Results were comparable when p-values were corrected for n=12 comparisons using the FDR method (p < .05 at time windows 250-300, 300-350, 350-400, 400-450, 450-500, and 500-600). Middle panel: averaged over all time windows and lateralities, separately for correct/error trials. Right panel: averaged over all time windows and correct/error trials, separately for ipsi/contralateral sensors (right panel).

A model that included the four CAPS symptom severity groupings separately revealed several significant interactions [Dysphoria severity and Congruency, Wald χ2(1) = 8.45, *p* = .003, Hyperarousal severity and Congruency, Wald χ2(1) = 5.33, *p* = .021], but post-hoc examination of these effects did not reveal any significant relationships between symptom domains and BBR. Overall, our analyses suggest that overall PTSD symptom severity may be associated with a failure to adaptively reduce motor inhibition to support response control, as indexed by increased peri-response BBR.

### 2.5 Error-Related Somatomotor Beta Burst Rates Are Associated with Overall PTSD Symptom Severity

The addition of diagnostic groups (controls/PTSD/mTBI/PTSD+mTBI) to our previous analysis of correct-error differences in BBR (incongruent trials only) demonstrated an interaction of diagnostic group by time window, Wald χ2(33) = 95.79, *p* < .001. Post-hoc examination of this effect found a difference between healthy controls and individuals with both mTBI and PTSD in a single time window, 50-100 ms (*z* = 2.67, *p* = .037), indicating higher pre-response BBR for healthy controls than for comorbid individuals. This contrast was no longer significant when results were corrected for n=12 time windows using the FDR method (Benjamini & Hochberg, 1995). When subjects with subthreshold PTSD were included in the PTSD groups, this analysis revealed an interaction of Group by Time Window, Wald χ2(33) = 48.82, *p* = .037, and an interaction between Group and Accuracy, Wald χ2(3) = 13.51, *p* = .007. However, post hoc interpretation of these interactions did not reveal any significant group differences.

A model that considered overall PTSD symptom severity (as assessed via the CAPS) rather than a formal diagnosis of PTSD showed a main effect of overall PTSD symptom severity, Wald χ2(1) = 4.54, *p* = .033, an interaction between overall PTSD symptom severity and Accuracy (correct/error), Wald χ2(1) = 11.56, *p* < .001, an interaction between overall PTSD symptom severity and Laterality, Wald χ2(1) = 5.18, *p* = .023, and an interaction between overall PTSD symptom severity and Time Window, Wald χ2(11) = 59.99, *p* < .001. Examination of the simple effect of CAPS on BBR within correct/error trials indicated that CAPS was significantly related to increased BBR in error trials (*z* = 2.81, *p* = .005) but not correct trials (*z* = 1.36, *p* > .1; *Figure 4B*). Examination of the simple effect of CAPS on BBR within contralateral/ipsilateral trials indicated that CAPS was significantly related to increased BBR contralateral to the flanker target stimulus (*z* = 2.57, *p* = .010) but not ipsilateral to the target stimulus (*z* = 1.60, *p* > .1; *Figure 4B*). Finally, examination of the simple effect of CAPS within each time window indicated that overall symptom severity was related to a diminished BBR reduction in peri-response time windows from 200-600 ms post-stimulus (200-250 ms *z* = 2.03, *p* = .042, 250-300 ms *z* = 2.62, *p* = .008, 300-350 ms *z* = 2.70, *p* = .007, 350-400 ms *z* = 2.70, *p* = .007, 400-450 ms *z* = 3.08, *p* = .002, 450-500 ms *z* = 3.07, *p* = .002, 500-550 ms *z* = 2.03, *p* = .043, 500-600 ms *z* = 2.47, *p* = .014; *Figure 4B*). These results remained significant for the 250-300, 300-350, 350-400, 400-450, 450-500, and 550-600 ms time windows (no longer significant at 200-250 and 500-550 ms) when *p*-values were corrected for n=12 multiple comparisons via the FDR (Benjamini & Hochberg, 1995) method.

A model that included CAPS symptom severity groupings revealed several significant interactions [Dysphoria severity and Accuracy, Wald χ2(1) = 8.82, *p* = .002, Avoidance severity and Accuracy, Wald χ2(1) = 12.69, *p* < .001, Intrusion severity and Accuracy, Wald χ2(1) = 17.80, *p* < .001], but post-hoc interpretation of these effects did not reveal any significant relationships between symptom domains and BBR. Our analyses suggest that overall PTSD symptom severity is associated with error-related BBR (rather than correct) and also appears specific to target-contralateral BBR (as opposed to distractor-contralateral BBR). Overall, PTSD symptom severity appears to be associated with a failure to adaptively reduce motor inhibition to support response control, as indexed by increased patterns of BBR.

### 2.6 Somatomotor Beta Burst Rates Mediate the Link Between Overall PTSD Symptom Severity and Impaired Conflict Monitoring

Given significant associations between overall PTSD symptom severity, peri-response BBR (250-550 ms), and accuracy rates, we tested the hypothesis that peri-response BBR might mediate the link between overall PTSD symptom severity and impaired conflict monitoring performance. We ran this analysis separately for correct congruent, correct incongruent, and error incongruent trial BBR and congruent/incongruent accuracy rates, respectively, for a total of three mediation analyses. For correct congruent trials, robust mediation analysis failed to demonstrate any significant indirect effects (*p* > .4). Robust mediation analysis also failed to demonstrate any mediating influence of error incongruent BBR (*p* > .9).

However, for correct incongruent trials, robust mediation analysis demonstrated a significant indirect effect of overall PTSD symptom severity on accuracy mediated by peri-response BBR, β: -.035, 95% bootstrap CI = [-.106, -.004], *p* = .027. Furthermore, prior to inclusion of BBR in the regression, overall PTSD symptom severity significantly predicted incongruent trial accuracy rates, β = -.19, *p* = .047. Following the addition of BBR to the model, overall PTSD symptom severity no longer predicted incongruent accuracy rates, β = -.13, *p* > .1. Thus, we conclude that correct incongruent BBR fully mediate the association between overall PTSD symptom severity and impaired conflict monitoring accuracy.

## 3 Discussion

### 3.1 General Discussion

We measured motor cortical indicators of cognitive control by characterizing beta-frequency bursts in a large sample of United States military veterans who had been deployed during either Operation Iraqi Freedom or Enduring Freedom. Beta burst rates (BBR) tracked motor control processes during conflict monitoring, correlated with individual differences in conflict monitoring, and were psychometrically reliable, suggesting that BBR are suitable for use as a measure of individual differences. Individual differences in peri-response BBR were related to dimensional posttraumatic stress symptom severity and explained (mediated) the influence of symptomatology on reduced conflict monitoring. Our investigation reveals that posttraumatic stress symptom severity modulates brain processing of cognitive control via inhibition at the motor representational level.

### 3.2 Beta Bursts Reveal the Dynamics of Motor Inhibition During Conflict Monitoring

Human somatomotor brain potentials in the beta frequency band reflect the output of cognitive control processes (Fischer et al., 2018; Soh et al., 2021). Physiologically, somatomotor beta-band activity is better characterized as an aperiodic burst-like process rather than a continuous and slow change in power (Shin et al., 2017). Somatomotor BBR decrease contralateral to the responding hand and predict individual differences in “go” response times (RTs) in the stop-signal task (Wessel, 2020), a common task examining response inhibition (Verbruggen et al., 2019). Furthermore, BBR increase precipitously following stop-signals (Wessel, 2020), suggesting BBR correspond to inhibition of inappropriate motor responses. We found that BBR were lower over contralateral (compared to ipsilateral) motor sensors for both congruent and incongruent correct trials, and ipsilateral, but not contralateral, BBR were lower for incongruent trials (compared to congruent). Since action preparation suppresses beta oscillations (Doyle et al., 2005; Pfurtscheller et al., 1996), BBR likely index disinhibition of ipsilateral motor cortex by distracting flanker stimuli. Correct/incorrect BBR did not differ contralateral to the target, that is, motor representations of target stimuli were equivalent for correct/error trials. However, error trials showed reduced BBR (compared to correct trials) over motor cortex contralateral to distractors. This result is undoubtedly confounded since individuals responded in the distractor direction on error trials, thus being at least partially driven by differences in the direction of response. Nevertheless, conflict monitoring errors appear driven by increased motor representations of distractor stimuli, rather than decreased motor representations of target stimuli.

Peri-response BBR negatively correlated with individual differences in accuracy rates for all task conditions, indicating that reduced peri-response motor disinhibition is associated with worse response control performance across subjects. Intriguingly, this effect was in the same direction over contralateral and ipsilateral sensors, where one might expect the correlation to be opposite in sign for ipsilateral sensors (i.e., increased suppression of distractor motor responses should improve accuracy). This is likely because BBR lateralization (contralateral<ipsilateral) is a within-subject effect, while BBR-accuracy correlations are a between-subjects effect. That is, lower contralateral (vs. ipsilateral) BBR reveals the mechanism of BBR to be inhibition of motor representations. Meanwhile, between subjects, increased BBR overall index increased motor inhibition.

While BBR appear sensitive to individual differences including Parkinson’s severity (Vinding et al., 2020), state anxiety (Sporn et al., 2020), and disorganization symptoms in psychosis (Briley et al., 2021), to our knowledge no report has yet characterized the *reliability* of beta burst rates. Given that reliable brain indices are required for characterizing individual differences and making clinical predictions (Button et al., 2013; Hedge et al., 2018; Zuo et al., 2019), we examined the split-half reliability of BBR in the flanker paradigm. Our approach assessed split-half reliability by bootstrapping random split-half averages per subject and correlating them across subjects, which avoids potential noise due to trials included in a single-split-half analysis (Macatee et al., 2021). Pre-correct-response BBR have good split-half reliability (Pearson r > 0.7) in the flanker paradigm, which increases to become very high in peri-response time windows (Pearson r > 0.8). Error-trial BBR were less psychometrically reliable, but still approached adequate levels during peri-response time periods (Pearson r > .65). Remarkably, this indicates that beta burst events show comparable split-half reliability to event-related potentials (ERPs) evoked during cognitive tasks. For example, the error-related negativity, an ERP component often used to assess individual differences in performance monitoring, has a split-half reliability estimated to be between 0.7 and 0.9 (Foti et al., 2013; Hajcak et al., 2019; Olvet & Hajcak, 2009a, 2009b; Riesel et al., 2013). Thus, beta burst rates carry potential as a measure of individual differences in neural responses subserving cognitive control.

### 3.3 Beta Burst Rates are Systematically Related to the Severity of Posttraumatic Stress

Our primary intention in the current report was to investigate if beta burst events could index response control dysfunction related to posttraumatic stress symptomatology in combat-exposed veterans. In line with prior reports of brain activation in combat-exposed veterans (Marquardt et al., 2021), we found that BBR predicted PTSD symptom severity but failed to be associated with a categorical diagnosis of PTSD. It is possible that the association with symptoms merely reflects an increase in statistical power with continuous as opposed to dichotomous measures (Altman & Royston, 2006; Lazic, 2008); however, the benefit of dimensional measures mirrors a recent focus on less reliance on diagnostic categories in psychopathology research. For example, the DSM-5 (2013) includes severity indicators for many disorders, consistent with the critical role that disorder severity plays in clinical management. The lack of an association of BBR with mTBI adds to the growing body of evidence that many reported consequences of military mTBI can be attributed to psychopathology other than mTBI, including PTSD (Disner et al., 2017; Marquardt et al., 2021).

Careful characterization of a large sample of individuals with varying levels of PTSD and mTBI enabled us to investigate influences of both common deployment-related conditions. Severity of posttraumatic stress, but not comorbid mTBI, was related to maladaptive motor cortex activity during conflict monitoring. These results were stronger for error trials than for correct trials, and for sensors contralateral to the flanker target stimulus (compared to ipsilateral). As such, increasing PTSD symptom severity was associated with impaired motor inhibitory processes related to target, rather than distractor, motor processing. We suggest that lower reactivity (reduction) in peri-response BBR could index difficulty in effectively reducing inhibitory drive in motor cortex and releasing the correct response to target stimuli. This is supported by the specificity of these BBR-symptom associations, which were strongest for incongruent target-contralateral and error conditions. The dysfunction in the somatomotor inhibitory mechanism explained (mediated) the link between increasing PTSD symptom severity and reduced response control, supporting the argument that dysfunctional somatomotor inhibition underlies conflict monitoring deficits in PTSD.

Future research should continue to elucidate other neural circuits affecting motor inhibitory processes. For example, in a stop-signal paradigm, prefrontal brain regions rapidly instantiate beta bursting at somatomotor sensors (Wessel, 2020), and inhibition can also be transmitted to motor cortex via beta bursting from the subthalamic nucleus that is transmitted to motor cortex via the thalamus (Diesburg et al., 2021). The effect of prefrontal beta bursting might be specific to outright response stopping, however, as we did not note any prefrontal increases in BBR in our topographic plots. While we did explore the possibility that conflict monitoring recruits prefrontal beta bursts the same way outright response stopping does (Wessel, 2020), we did not find any congruency effects on BBR at prefrontal sensors. Somatomotor beta bursts, the motor inhibitory mechanism underlying PTSD-relevant cognitive control deficits, might also be under the control of executive brain regions via cross-frequency interactions (for example frontal midline theta). In healthy controls, but not individuals with PTSD, somatomotor beta activity was linked to prefrontal theta-band activation (Cohen et al., 2013) – a common substrate underlying multiple types of cognitive control (Cavanagh et al., 2012; Cavanagh & Frank, 2014; Eisma et al., 2021). Furthermore, in a sample of trauma-exposed veterans, increased cognitive control-related brain activation in the anterior cingulate cortex (ACC), a key brain region regulating cognitive control computations (Botvinick, 2007; Botvinick et al., 2004; Gehring & Knight, 2000; Sohn et al., 2007), was associated with reduced PTSD symptom severity (White et al., 2018). This suggests PTSD is characterized by a failure of top-down motor modulation paired with dysregulated motor output systems. Thus, PTSD-related impairments in executive and somatomotor brain regions might be linked.

### 3.4 Limitations

A limitation of the current report is the cross-sectional nature of the military sample. Since soldiers were recruited and studied following deployment, we are unable to assess whether the observed associations represent a consequence of posttraumatic stress or a risk factor for developing more severe posttraumatic stress. Examinations that measure recruits prior to and following trauma exposure are needed to disambiguate these possibilities. Such studies could also allow the potential development of adaptive reduction of motor inhibition into a mature biomarker for resilience to PTSD following trauma exposure. Additionally, PTSD is commonly associated with psychopathological comorbidities including anxiety, depression, and alcohol use disorder, all of which are associated with altered cognitive control (Bogg et al., 2012; Cavanagh et al., 2017; Grahek et al., 2019; Krug & Carter, 2010; Rawls et al., 2021; Zilverstand et al., 2018). While control analyses demonstrated that none of the observed motor inhibition or conflict monitoring effects were confounded by these diagnoses (Generalized Anxiety/Major Depression/Alcohol Dependence), deeper consideration of comorbidities is essential to understand the complex clinical presentation of PTSD. Future studies might address this deficit by oversampling individuals with specific comorbidities. With thorough and dimensional phenotyping, it is possible that this approach might reveal BBR as a transdiagnostic neural mechanism providing a readout of impaired conflict monitoring that spans diagnostic labels. Finally, as in prior examinations of beta-bursting in scalp EEG, we recorded a relatively low proportion of trials containing beta-bursts (in some time windows, only 4-20% of trials contained a beta burst). While this could suggest that beta bursts are not frequent enough to completely explain motor inhibition (Errington et al., 2020), this could also be due to the relatively low signal-to-noise ratio of beta activity in scalp EEG (Wessel, 2020), and beta burst counts are also sensitive to the threshold used to define bursts (Shin et al., 2017). As such, while we recorded relatively low BBR during this task, these rates are in line with published estimates of BBR and showed very good reliability despite the low incidence of bursts.

### 3.5 Conclusion and Future Directions

Our analysis of the role of somatomotor beta-burst events demonstrates that beta bursts index critical brain mechanisms of cognitive control and reveal failures to adaptively reduce motor inhibition in PTSD. Findings of the current study support the possibility of using training of response control to improve aspects of PTSD symptomatology. Furthermore, since beta bursts occur in single trials of EEG, we suggest a possible focus on noninvasive closed-loop neuromodulation of human somatomotor cortex triggered by real-time detection of beta burst events to detect and correct conflict monitoring failures in real time. Such a stimulation protocol could be titrated to an individual’s average beta-frequency power during a calibration session, and excitatory neuromodulation could be used to adaptively disinhibit motor cortex during periods of pathologically high beta bursting. Such a stimulation protocol would clarify the causal role of somatomotor BBR interventions in support of cognitive control functions. This protocol would also allow testing whether a neuromodulatory intervention at the level of the motor cortex could generalize to brain regions responsible for non-motor functions, for example emotion regulation. It appears that consideration of dimensional aspects of posttraumatic psychopathology is critical to understanding and eventually treating the neural basis of trauma-related dysfunction.

## 4 Materials & Methods

### 4.1 Participants

The sample was n = 130 US military veterans (see *Table 1* for demographics) recruited through the Minneapolis VA Health Care System (VAHCS). Participants were recruited based on their history of deployment to combat zones in either Iraq and/or Afghanistan (Operations Iraqi Freedom and Enduring Freedom, respectively). Recruitment targeted veterans with likely posttraumatic stress disorder (PTSD) diagnoses as well as non-treatment-seeking veterans with similar deployment experiences for recruitment. Study procedures were approved by the Institutional Review Boards at the VAHCS and the University of Minnesota, and study participants gave written informed consent prior to undergoing the study procedures. This sample has been previously reported in (Davenport et al., 2016; Disner et al., 2018; Marquardt et al., 2021); none of these publications have examined data from the flanker paradigm. Error trial analysis considered a smaller subset of these participants (n = 106) who had at least ten error trials with clean EEG (all n = 130 subjects had sufficient correct trial data).

**Table 1.** Demographic and clinical characteristics of sample.

| Variable | No PTSD |  |  |  |  |  | PTSD+Subthreshold |  |  |  |  |  |
| --- | --- | --- | --- | --- | --- | --- | --- | --- | --- | --- | --- | --- |
|  | No mTBI |  |  | mTBI |  |  | No mTBI |  |  | mTBI |  |  |
|  | *n* | *M* | *SD* | *n* | *M* | *SD* | *n* | *M* | *SD* | *n* | *M* | *SD* |
| Total Count | 61 |  |  | 25 |  |  | 22 |  |  | 22 |  |  |
| Female | 3 |  |  | 0 |  |  | 1 |  |  | 0 |  |  |
| Age (years) |  | 34.78 | 8.49 |  | 32.05 | 8.62 |  | 36.18 | 7.97 |  | 33.17 | 6.08 |
| Education (years) |  | 14.83 | 1.56 |  | 14.14 | 1.67 |  | 14.82 | 2.22 |  | 14.06 | 2.01 |
| Depressive Disorder<br>Diagnosis | 8 |  |  | 2 |  |  | 4 |  |  | 10 |  |  |
| Alcohol Dependence<br>Diagnosis | 0 |  |  | 0 |  |  | 2 |  |  | 2 |  |  |
| CAPS Severity (total) |  | 20.5 | 15.7 |  | 18.4 | 12.3 |  | 50 | 28.6 |  | 55.5 | 23.9 |
| CAPS Intrusive<br>Reexperiencing Total |  | 4.2 | 4.96 |  | 5.36 | 6.7 |  | 13.1 | 8.88 |  | 16.6 | 7.44 |
| CAPS Avoidance Total |  | 1.66 | 2.78 |  | 0.72 | 1.4 |  | 6.32 | 4.81 |  | 6.82 | 3.95 |
| CAPS Dysphoria Total |  | 10.2 | 8.96 |  | 7.04 | 6.09 |  | 22.5 | 12.9 |  | 23.6 | 12.6 |
| CAPS Hyperarousal<br>Total |  | 4.44 | 3.39 |  | 5.24 | 3.06 |  | 8.09 | 4.18 |  | 8.45 | 3.76 |
| MN-BEST Blast mTBI<br>Severity |  | 0 | 0 |  | 3.04 | 1.80 |  | 0 | 0 |  | 3.23 | 1.35 |

PTSD = posttraumatic stress disorder, mTBI = mild traumatic brain injury, *N* = count, *M* = mean, *SD* = standard deviation. “+Subthreshold” reflects individuals who meet criteria for at least one symptom from each symptom domain of DSM-IV PTSD.

### 4.2 Clinical Assessment

We conducted interview-based assessments for psychopathology using the Structured Clinical Interview for DSM-IV Axis I Disorders excluding the PTSD module (SCID-I; (First & Gibbon, 2004)). We assessed for posttraumatic stress using the Clinician-Administered PTSD Scale for DSM-IV [CAPS, fourth edition; (Blake et al., 1995; Weathers et al., 2001)]. We achieved consensus for categorical psychiatric diagnoses with assessment teams that reviewed all available research and clinical information and included at least one licensed doctoral-level clinical psychologist. We also determined the PTSD and “subthreshold” PTSD diagnoses using the same consensus procedure. Individuals were labeled as having subthreshold PTSD if they endorsed at least one symptom in each DSM-IV-TR Criterion B-D symptom groupings and met the threshold for being exposed to a traumatic event (Criterion A). We subdivided the CAPS into four subscales based on previous analyses of the factor structure of DSM-IV(-TR)-defined posttraumatic stress symptomatology (Palmieri et al., 2007; Simms et al., 2002; Yufik & Simms, 2010). By summing the CAPS frequency and intensity scores, we produced dimensional measures of symptom severity within the domains of intrusive reexperiencing (B1-B5), avoidance (C1, C2), dysphoria (C3-D3), and hyperarousal (D4, D5). For our analyses using the CAPS, we first examined overall severity score (all factors summed) then followed up with additional analyses using the four symptom groupings.

We assessed participant history of likely mTBI events using the semi-structured Minnesota Blast Exposure Screening Tool (MN-BEST; Nelson et al., 2011), focusing on the three most severe self-identified deployment-related blast exposure events. We achieved consensus on mTBI via assessment teams that included at least one licensed clinical neuropsychologist (Nelson et al., 2015). Blast-induced mTBI severity was quantified using an adaptation of the (Ruff & Richardson, 1999) rating scheme.

### 4.3 Conflict Monitoring Task

Participants completed an Eriksen Flanker paradigm that presented congruent (<<<<< or >>>>>) or incongruent (<<><< or >><>>) sets of arrows (50% incongruent). Participants were instructed to respond with their left thumb if the middle arrow pointed left, and to respond with their right thumb if the middle arrow pointed right, while ignoring the distractor arrows on either side of the target arrow. Participants completed 400 trials, which were divided into 4 blocks with self-paced breaks in between. The task required approximately 20 minutes to complete.

### 4.4 EEG Acquisition and Preprocessing

EEG was sampled at 1024 Hz using a 128-channel BioSemi ActiveTwo EEG system, acquired reference-free (via CMS/DRL sensors) and re-referenced to Cz upon import. Data were automatically preprocessed using EEGLAB 2021 (Delorme & Makeig, 2004) and MATLAB R2021a. Continuous data were high-pass filtered at 0.5 Hz (transition band width of 0.5 Hz) and low-pass filtered at 35 Hz (transition band width of 10 Hz) using zero-phase least-squares FIR filters, then downsampled to 250 Hz. Bad channels were detected using joint probabilities (3 SD cutoff) and removed. Data were epoched around the flanker arrow stimulus (400 ms before to 700 ms after) and trials containing non-stereotyped artifacts were removed using a cutoff of ±500 μV prior to ICA computation. Temporal infomax ICA (Makeig et al., 1996) was computed, and components capturing stereotyped artifacts were removed using ICLabel (Pion-Tonachini et al., 2019). Trials containing residual artifacts were detected and removed using a voltage cutoff of ±125 μV. Single trials with improbably fast RTs (<100 ms) were removed from the EEG, as were trials where the participant did not respond. Deleted channels were spherically interpolated, channel Cz was added back to the data, and single trials of artifact-free EEG were re-referenced to the montage average. To sharpen the focal motor topographies of beta bursts (Wessel, 2020) and to improve our ability to resolve lateralized brain responses by removing volume-conducted activity (Kayser & Tenke, 2015), we applied the surface Laplacian transform using the CSD Toolbox (https://psychophysiology.cpmc.columbia.edu/software/csdtoolbox/index.html) with default parameters. This transformed the EEG to a reference-free current scalp density representation, which we employed for all further analyses.

### 4.5 Quantification of Beta Bursts from Scalp EEG

Beta burst detection was performed exactly as described by (Shin et al., 2017) and (Wessel, 2020); the following description is adapted from those works. Single trials of artifact-free EEG were transformed to a time-frequency representation by convolving the raw data with a family of Morlet wavelets described by

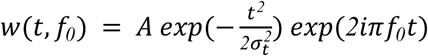

for each frequency of interest *f*_*0*_, where 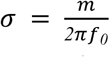, normalization factor 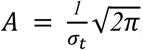, and *m*=*7* cycles for each of 15 evenly spaced frequencies spanning the beta band (15-29 Hz). Time-frequency power estimates were calculated by squaring the magnitude of the complex result of the convolution. All within-subject time-frequency power estimates were normalized by frequency-specific medians calculated across all time points (i.e., -400 to 700 ms surrounding stimulus onset) and all single trials (Shin et al., 2017). Following this power normalization, which corrected both for differences in average beta power between subjects and between frequencies, beta bursts in single trials were defined as local maxima in the time-frequency surface (using the MATLAB function *imregionalmax*) that exceeded an *a priori* threshold of 6 times the median power (Shin et al., 2017; Wessel, 2020).

### 4.6 Data Reduction for Beta Burst Rate Analysis

We defined 12 50-ms time bins extending from the onset of the flanker stimulus to 600 ms post-stimulus (0-50 ms, 50-100 ms, 100-150 ms, 150-200 ms, 200-250 ms, 250-300 ms, 300-350 ms, 350-400 ms, 400-450 ms, 450-500, 500-550 ms, 550-600 ms). Time windows were defined inclusively, so that a sample at 48 ms went in the 0-50 ms bin and a sample at 52 ms went in the 50-100 ms bin. For samples that landed exactly on a start/end point of a time window (e.g., 100 ms), the sample was included in both time windows. Since the average response time for correct incongruent trials was 501 ms, this measurement window ensured that we captured beta bursting occurring between stimulus presentation and response for most trials. Within each time bin, we counted the number of bursts that occurred in each single trial and for each sensor. These beta burst counts were averaged over all trials, forming our primary dependent variable “beta burst rate” (BBR). As in prior applications of this method, BBR are thus measured as n(beta-bursts)/n(trials), where each window typically has either 0 or 1 (but sometimes more) beta burst(s). BBR measured following this procedure are interpreted as the average number of beta bursts in each time window per trial. BBR were extracted from small spatial clusters of lateral somatomotor sensors surrounding C3 (left) and C4 (right), and separately averaged (over trials) into four beta burst time series according to factors of congruency (congruent vs. incongruent) and laterality (ipsilateral vs. contralateral). For incongruent trials only, we also averaged beta bursts for incorrect trials. Together, these procedures produced our primary dependent variable “beta burst rate,” (BBR) or the average number of beta bursts that occur each trial (per time window and trial type).

### 4.7 Reliability Analysis of Beta Burst Rates

While recent studies have begun examining the relationships of BBR with individual difference factors [e.g., RT relationships in (Wessel, 2020), Parkinsonian disease severity in (Vinding et al., 2020)], none of these recent examinations have considered the important point of whether their investigation provided a *reliable* measure of BBR. Psychometric reliability is critical for interpretation of brain-disease relationships and for development of robust biomarkers of disease (Button et al., 2013; Hedge et al., 2018; Zuo et al., 2019). We characterized the reliability of our measured BBR, separately for congruent (incongruent) trials and ipsi(contra)-lateral sensors. We estimated the average split-half reliability, as well as 95% confidence intervals (CIs) surrounding the estimate, by generating random split-half across-trial averages per subject (5000 repetitions) and correlating each of the split-half estimates together across subjects (Pearson correlation, corrected for halving the sample size using Spearman-Brown correction). Note that while BBR deviated from normality and Pearson correlations are sensitive to deviations from normality (Kowalski, 1972), we used Pearson correlations for this analysis to render our reliability metrics comparable to the existing EEG reliability literature (Luking et al., 2017; Olvet & Hajcak, 2009b, 2009b; Riesel et al., 2013; Towers & Allen, 2009).

### 4.8 Beta Burst Rate and Task Performance Associations

We tested the hypothesis that BBR index trait-like individual differences in cognitive control operations. For all considered variables, we first checked the normality of data distributions. BBR, RT, and accuracy rates were all visibly non-normal, which was verified by significant Shapiro-Wilk tests for all variables. As such, we used Kendall’s tau rather than Pearson correlations because the distribution of Pearson’s r is not robust to non-normality (Kowalski, 1972), and Kendall’s tau has lower gross error sensitivity (i.e., is more robust) than Spearman’s correlation (Croux & Dehon, 2010). We calculated bivariate correlations between accuracy rates and trial-averaged BBR, and between response times and trial-averaged BBR, separately for ipsilateral and contralateral sensors and for congruent and incongruent trials, as well as for error trials (incongruent only). We constructed 95% bias-corrected accelerated bootstrap CIs (5000 repetitions) surrounding each of these correlation coefficients using the ‘confintr’ package, version 0.1.2 (Mayer, 2022).

### 4.9 Statistical Analysis of Behavior and Beta Burst Rates

For statistical analyses of behavioral responses (accuracy, RT) and BBR, we fit linear mixed-effect models (LMMs) using the ‘lme4’ package, version 1.1-29 (Bates et al., 2015, 2022). LMMs are robust against violations of distributional assumptions (Schielzeth et al., 2020), being robust for data with moderate skewness and even extreme kurtosis with sample sizes > 60 (Arnau et al., 2013). Of our primary outcome variables, only accuracy rates exhibited concerning skewness (skewness = -1.62; all other outcome variable |skewness| < 0.68). As such, for LMMs predicting accuracy rates, we compared the results of LMMs fit using lme4 to those fit using the ‘robustlmm’ package, version 3.0-4 (Koller, 2016) to ensure that resulting estimates replicated using robust model estimation. We estimated robust LMM *p*-values using model *t*-statistics and Satterthwaite-approximated degrees-of-freedom as in (Geniole et al., 2019).

LMMs predicting accuracy rates and response times used a single within-subject factor of Congruency (congruent/incongruent), and LMMs predicting correct-trial BBR used within-subject factors of Congruency (congruent/incongruent), sensor Laterality (ipsilateral/contralateral), and Time Window (50 ms levels). Note that sensor Laterality refers to sensor location relative to the direction of the target stimulus (center arrow). All LMMs examining error-related BBR instead included within-subject factors of Outcome (correct/error), sensor Laterality (ipsilateral/contralateral), and Time Window (50 ms levels), for incongruent trials only (since subjects made almost no errors on congruent trials). In analyses comparing correct and error trials, “contralateral” refers to sensors on the side opposite the target stimulus and “ipsilateral” refers to sensors on the same side as the target stimulus (i.e., contralateral to the distractor stimuli). As such, this analysis directly compares motor inhibitory processing of target and distractor stimuli.

Models included one or more between-subjects factors describing clinical presentation. First, we ran models with a between-subjects factor of diagnostic group (four levels: controls, PTSD, mTBI, both PTSD and mTBI). We used two versions of this diagnosis-based model; the second enlarged the PTSD groups to include participants with subthreshold, but clinically significant, PTSD symptomatology. We then analyzed models that used overall PTSD symptom severity (CAPS; z-scored across participants), to examine whether increasing severity of PTSD symptomatology predicted cognitive control performance or BBR. Finally, we considered models that used each of the four CAPS subscales (intrusion, avoidance, dysphoria, hyperarousal; z-scored across participants) as the primary between-subjects factors. All models contained a random intercept per participant, and all severity-based models covaried for mTBI severity. All models included interaction terms between all within-subjects factors and the primary between-subjects factors.

Significance of individual factors was assessed using type III Wald chi-square tests implemented with the R ‘car’ package, version 3.0-13 (Fox et al., 2022). Significant categorical-categorical interactions were characterized using the emmeans function, and significant categorical-continuous interactions were assessed by comparing trends within each categorical factor level using the emtrends function, both implemented in the R ‘emmeans’ package, version 1.7.4-1 (Lenth et al., 2022). For post hoc testing of significant BBR interactions at each level of Time Window, we present results both without further correction (i.e., Fisher’s Least Significant Difference) and following correction for n=12 multiple comparisons using the false discovery rate (FDR) method with *q* < .05 taken as evidence of a significant result (Benjamini & Hochberg, 1995).

### 4.10 Mediation Analysis

Given the significant associations between overall PTSD symptom severity, peri-response BBR, and accuracy rates,we ran follow-up analyses to examine whether peri-response BBR (averaged over time windows comprising 250-550 ms post-stimulus) mediated the link between overall PTSD symptom severity and accuracy, separately for correct congruent, correct incongruent, and incorrect incongruent trials. Often ordinary least squares (OLS) regression is used for mediation analysis, with CIs for the indirect (causal mediation) effect estimated via parametric bootstrapping. However, the parametric bootstrap for indirect effects fits a mediation model to the data using normal-theory Maximum Likelihood (ML) estimation (Tofighi, 2020), which is easily distorted by even small deviations from normality. As noted previously, several measures were non-normal in our sample, suggesting that OLS mediation is not a proper approach for this analysis. Instead, we conducted outlier- and distribution-robust mediation analysis (Alfons et al., 2022a) using the R ‘robmed’ package, version 1.0.0 (Alfons et al., 2022b). This test used robust regression with 95% CIs estimated via percentile bootstrapping (5000 repetitions). We used the total CAPS score as the independent variable, accuracy rates as the outcome variable, and peri-response BBR (250-550 ms) as the mediating variable. Causal mediation analyses covaried for potential confounding effects of mTBI severity. Variables were z-scored prior to model fitting.

## CRediT Author Statement

ER: Conceptualization, Methodology, Software, Validation, Formal Analysis, Investigation, Writing - Original Draft, Writing - Review and Editing, Visualization. CAM: Validation, Data Curation, Writing - Review and Editing. SRS: Resources, Data Curation, Writing - Review and Editing, Supervision, Project Administration, Funding Acquisition.

## Acknowledgments

ER is supported by the National Institutes of Health’s National Center for Advancing Translational Sciences, grants TL1TR002493 and UL1TR002494. This work was funded through a Department of Veterans Affairs Rehabilitation R&D Program (I01RX000622) grant to SRS. In addition, this project was supported with resources and the use of facilities at the Minneapolis VA Health Care System. The content is solely the responsibility of the authors and does not necessarily represent the official views or policy of the U.S. Department of Veterans Affairs, the United States Government, or the National Institutes of Health’s National Center for Advancing Translational Sciences. We acknowledge code graciously hosted by Stephanie Jones (https://github.com/jonescompneurolab/SpectralEvents) and Jan Wessel (https://osf.io/v3a78/) for analyzing beta bursts.

The authors have no conflicts of interest to report.

